# Self-supervised Internal Learning Enhances Isotropic Resolution for Three-dimensional Fluorescence Microscopy

**DOI:** 10.64898/2026.06.04.717237

**Authors:** Mingzhe Wei, Pengcheng Xu, Junyu Liu, Xuesong Li, Xuhui Feng, Jun Zhu, Renwei Dong, Hengjia Ran, Wentao Zhu, Yubing Han, Yue Li, Min Guo, Huafeng Liu

## Abstract

Three-dimensional fluorescence microscopy often exhibits anisotropic resolution because axial information is poorly sampled and more blurred than lateral information, which complicates quantitative interpretation of fine 3D structures. Although optical remedies and computational restoration have been explored, many approaches require demanding system calibration or rely on accurate PSF models and assumptions that are difficult to satisfy across all samples and modalities. Here we present DeepIso, a self-supervised isotropy restoration framework that couples supervised pretraining with an internal-learning inference stage to estimate degradation directly from the measured volume. Without explicit PSF specification or enforced lateral–axial structural equivalence, DeepIso recovers axial frequency content and improves the continuity of elongated structures while retaining fine features, with superior performance over existing computational approaches in terms of both visual inspection and quantitative metrics. The method is validated on synthetic benchmarks and experimental datasets, demonstrating isotropy enhancement across confocal, light-sheet, and 3D structured illumination microscopy, thereby supporting downstream volumetric analysis including segmentation and tracking.

## 1. Introduction

Fluorescence microscopy is an essential tool in life sciences, providing high-resolution and high-specificity imaging vital for investigating cellular structures^1^, cell interactions^2^, disease mechanisms^3^, and drug development^4^. Unlike two-dimensional (2D) imaging, which captures a single plane, three-dimensional (3D) fluorescence imaging reconstructs the specimen in its entirety, offering more comprehensive volumetric coverage of biological sampless^5–7^. However, 3D imaging via z-stack acquisition poses inherent challenges. Due to the intrinsic diffraction limit of optics and larger axial sampling intervals, single image volumes exhibit pronounced anisotropic resolution – axial information is resolved more poorly than lateral information. This anisotropy significantly hinders accurate dissection and quantitative analysis of intricate 3D biological structures.

To enhance biological interpretations, researchers have proposed various strategies aimed at achieving isotropic or near-isotropic resolution in 3D fluorescence imaging. Hardware-based strategies have emerged, modifying optical systems to extend axial information through interference-based approaches^8–11^, multiview acquisitions^12–15^, and axially swept light-sheet microscopy (ASLM)^16,17^. Although effective, these methods often impose practical constraints, such as stringent alignment requirements, increased optical complexity, and the need for precise calibration^18–21^. Furthermore, the associated reconstructions frequently require computationally intensive workflows for multiview fusion, hindering widespread adoption ^22^.

Alternatively, computational image restoration methods^23–31^ offer another pathway toward resolution isotropy. For instance, 3D deconvolution algorithms reassign out-of-focus light, thereby improving the recovery of axial information. Nonetheless, the intrinsic physical and sampling constraints still lead to a narrower axial optical transfer function (OTF) compared to its lateral counterpart, limiting the recovery of high-frequency axial information. To overcome this limitation, deep learning-based restoration techniques have been developed, effectively extracting structural features from the data. Existing supervised methods, such as content aware image restoration (CARE)^23^, rely on accurate estimates of the axial point spread function (PSF) and the associated sampling model, which can lead to inaccuracies and mismatched reconstructions if estimations are not precise. By contrast, unsupervised generative adversarial network (GAN) approaches, including CycleGAN variants^32^ and SelfNet^33^ frameworks, are typically trained on unpaired lateral and axial datasets. Their success hinges on the implicit assumption of structural similarity between lateral and axial views to establish meaningful cross-view mappings. Adversarial training, however, frequently suffers from optimization instability, making it highly sensitive to hyperparameters and run-to-run variability, which heightens the chance of producing artifacts or hallucinated textures. More recently, the SSAI-3D framework^34^ reformulates isotropy restoration as semi-blind axial deblurring by transferring knowledge from a natural-image-pretrained deblurring backbone to microscopy data through sparse fine-tuning. While this approach shows promise, it does have limitations, including incomplete axial resolution recovery and potential complications in reconstructing high-resolution images due to the fundamental degradation model mismatch between microscopy images and natural images.

Building on this body of prior work, we introduce DeepIso, a self-supervised framework designed to enhance isotropic resolution in fluorescence microscopy through internal learning. DeepIso integrates supervised pretraining with a self-adaptive inference stage, enabling the system to learn degradation characteristics directly from the data without requiring accurate PSF estimation or assuming structural similarity between lateral and axial views. Unlike existing methods, DeepIso expands the effective axial-frequency bandwidth and recovers complex axial morphology with high fidelity, preserving continuity of elongated structures and detailed features. We validated the performance and reconstruction fidelity of DeepIso on both synthetic benchmarks and experimental datasets, demonstrating robust isotropy restoration across multiple microscopy modalities, including confocal, light-sheet, and 3D structured illumination microscopy. In all cases, DeepIso achieves superior performance over existing computational methods. DeepIso provides a practical solution for sample-adaptive isotropy enhancement, with broad potential for downstream analyses in volumetric visualization, segmentation, tracking, and long-term time-lapse analysis in diverse biological applications.

## 2. Results

### 2.1. Framework of DeepIso for isotropic resolution enhancement

To achieve isotropic restoration from inherently anisotropic microscopy data, we developed DeepIso, a self-supervised two-stage framework designed to bridge the resolution gap between lateral and axial dimensions (**Fig. 1**).

**Fig. 1.**
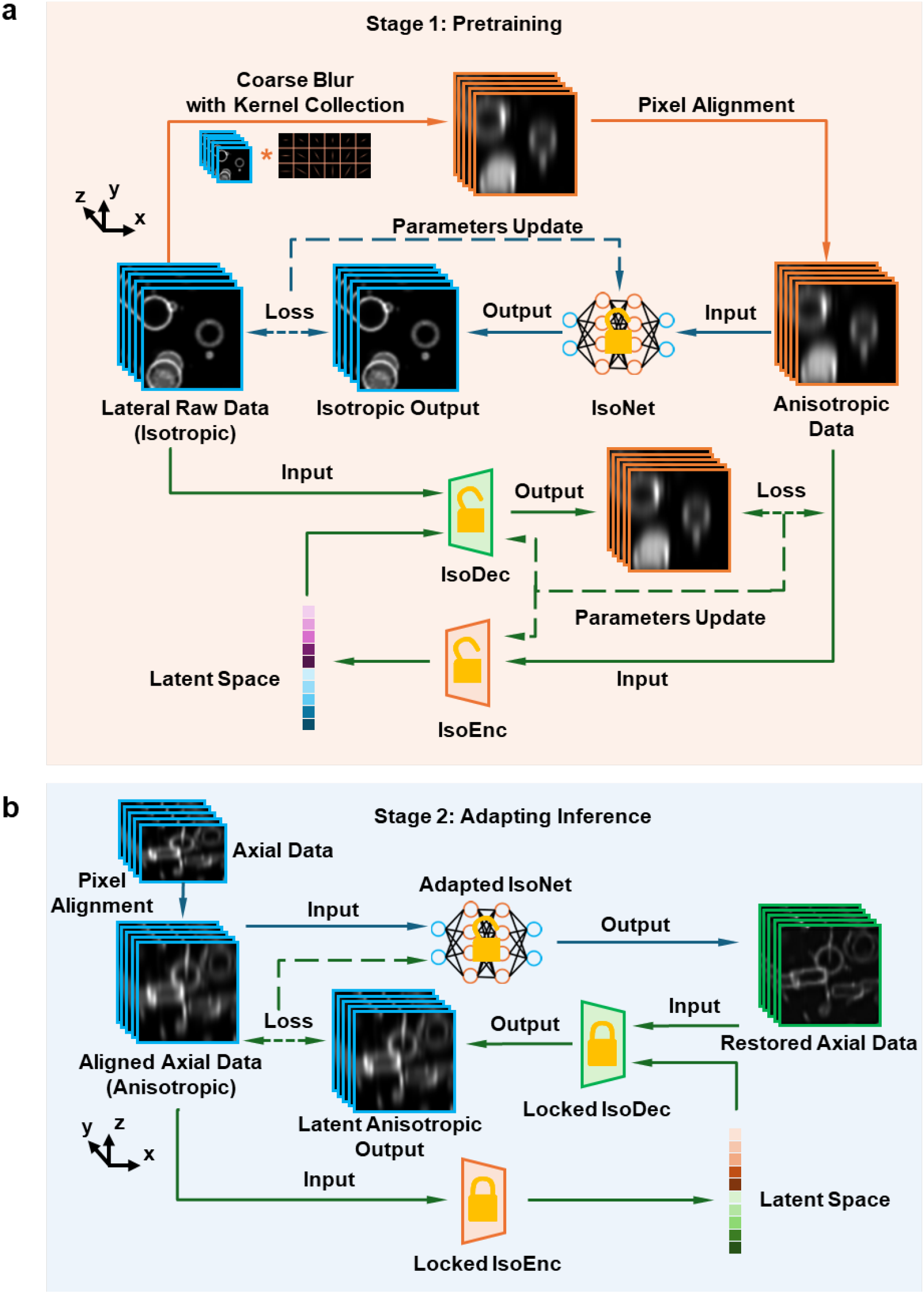
The two-stage DeepIso framework for self-supervised isotropic resolution restoration. DeepIso transfers knowledge from diverse, synthetically generated degradation patterns to the anisotropic restoration of specific dataset through a pretraining-then-adaptation workflow. **a**, Stage 1 (Pretraining): Supervised pretraining on synthetic data. An IsoNet backbone is trained to learn isotropic representations from artificially degraded data. Isotropic lateral slices are synthetically degraded using a collection of coarse axial blur kernels and pixel size alignment to generate diverse and realistic axial anisotropic candidates (Anisotropic Data). Then these axial anisotropic slices are fed into the IsoNet to learn the anisotropic-to-isotropic mapping by supervised pretraining. Meanwhile, an encoder module (IsoEnc) is concatenated to capture anisotropic structural features from anisotropic slices in the latent space, while a decoder module (IsoDec) is trained to utilize latent features to degrade the isotropic lateral slices by enforcing reconstruction consistency. IsoEnc and IsoDec are jointly updated to realize accurate latent feature representation. **b**, Stage 2 (Adapting Inference): The raw axial data (Axial Data) is first axially interpolated to match the axial pixel size to the lateral pixel size (Pixel Alignment) and get the aligned axial data for network input (Aligned Axial Data). These data are fed into IsoNet using transfer learning to get initial estimates (Restored Axial Data). Then these initial estimates together with the aligned axial data are used to generate the intermediate anisotropic outputs (Latent Anisotropic Output) using locked IsoEnc/IsoDec pair. By optimizing the consistency between the real axial data (Aligned Axial Data) and intermediate outputs (Latent Anisotropic Output), the IsoNet is iteratively updated and produces the final restored data (Restored Axial Data).

In Stage-1 Pretraining (**Fig. 1a**), an IsoNet backbone is trained to learn isotropic representations from synthetically degraded data. To emulate diverse optical conditions and axial blurring, we construct a coarse blur kernel library consisting of one-dimensional kernels of multiple orientations and scales (**Methods**). These kernels are convolved with isotropic lateral slices to generate pseudo-anisotropic inputs. Before training, a pixel alignment step is applied by downsampling one spatial axis to match the axial pixel size and subsequently upsampling it back to the lateral pixel size, ensuring uniform voxel spacing and consistent receptive-field sampling across all directions. The IsoNet is then optimized by minimizing the reconstruction loss between the isotropic ground truth and its blurred counterpart, yielding a pretrained model that captures scale-invariant and orientation-invariant isotropic features. In parallel with IsoNet pretraining, we introduce an auxiliary encoder–decoder pair (IsoEnc/IsoDec) to explicitly learn a compact latent representation of the anisotropic observation manifold. Specifically, IsoEnc encodes the pseudo-anisotropic volumes generated by the coarse-kernel collection into a low-dimensional latent space, and IsoDec is trained to reconstruct the corresponding anisotropic volumes by minimizing a self-reconstruction loss. This bottlenecked latent embedding regularizes the learning of isotropic features by enforcing cross-domain consistency: the restored isotropic prediction should remain explainable under the learned anisotropic generative pathway, rather than relying on unconstrained hallucination. The overall design is motivated by developments in internal learning and self-/zero-shot super-resolution, where supervised pretraining is complemented by test-time or sample-specific adaptation to bridge the gap between synthetic degradations and real acquisitions (e.g., ZSSR^35,36^, meta-transfer^37^, and related self-supervised fine-tuning strategies^38^). Consequently, Stage-1 establishes a stable, modality-agnostic latent space that can be locked in the following inference stage to provide a reliable internal constraint during test-time adaptation.

Importantly, unlike many GAN-based isotropic restoration methods, DeepIso does not rely on the implicit assumption that biological structures should appear similar between lateral and axial views. In Stage-2 Adapting Inference (**Fig. 1b, Supplementary Fig. 1, Supplementary Fig. 2**), the pretrained network is internally adapted to real experimental data exhibiting anisotropic sampling. During this process, the encoder–decoder pair (IsoEnc/IsoDec) is locked to preserve the latent-space consistency learned from pretraining, while only the intermediate IsoNet parameters are fine-tuned to accommodate the domain shift. This internal adaption permits inference on raw anisotropic volumes without external training data or explicit supervision, and it does so without invoking the common assumption of structural similarity between lateral and axial views. Consequently, DeepIso yields isotropic restorations that preserve axial morphology with high fidelity while suppressing spurious artifacts across imaging views.

### 2.2. Class-leading performance of DeepIso on simulated volumetric data

We first validated the isotropic restoration capability of DeepIso using simulated volumetric data, where both anisotropic inputs and isotropic ground truths were available for direct quantitative comparison (**Fig. 2**). Anisotropic inputs were obtained by blurring the simulated ground truth with a simulated point spread function (PSF) to mimic the anisotropic diffraction-limited response set by the objective lens and sampling. The simulation volume comprised a heterogeneous mixture of primitives, including point emitters, line segments, hollow spheres, and hollow cylinders with varying sizes, positions, and 3D orientations. Notably, for hollow cylinders, the circular end-cap is predominantly visible in lateral views, whereas the cylindrical sidewall becomes apparent only in axial views. This design intentionally breaks lateral–axial structural similarity, emulating specimens where cross-view appearance differs substantially. After reconstruction, DeepIso effectively recovered isotropic structures across the entire 3D volume, yielding results that closely matched the ground-truth references (**Fig. 2a**). Detailed inspection of representative sub-regions (**Fig. 2b–e**) demonstrated that DeepIso markedly improved axial views compared with existing deep-learning-based restoration models, including CARE^23^, CycleGAN^32^, SelfNet^33^, and SSAI3D^34^. DeepIso preserved membrane continuity (yellow arrows), resolved fine tubular and vesicular features (blue and green arrows), and minimized artifacts such as stripe-like hallucinations that were frequently observed in CycleGAN and SelfNet outputs. The recovered structures exhibited smooth intensity transitions along the axial direction, confirming the model’s ability to learn depth-aware isotropic representations rather than planar deconvolution priors.

**Fig. 2.**
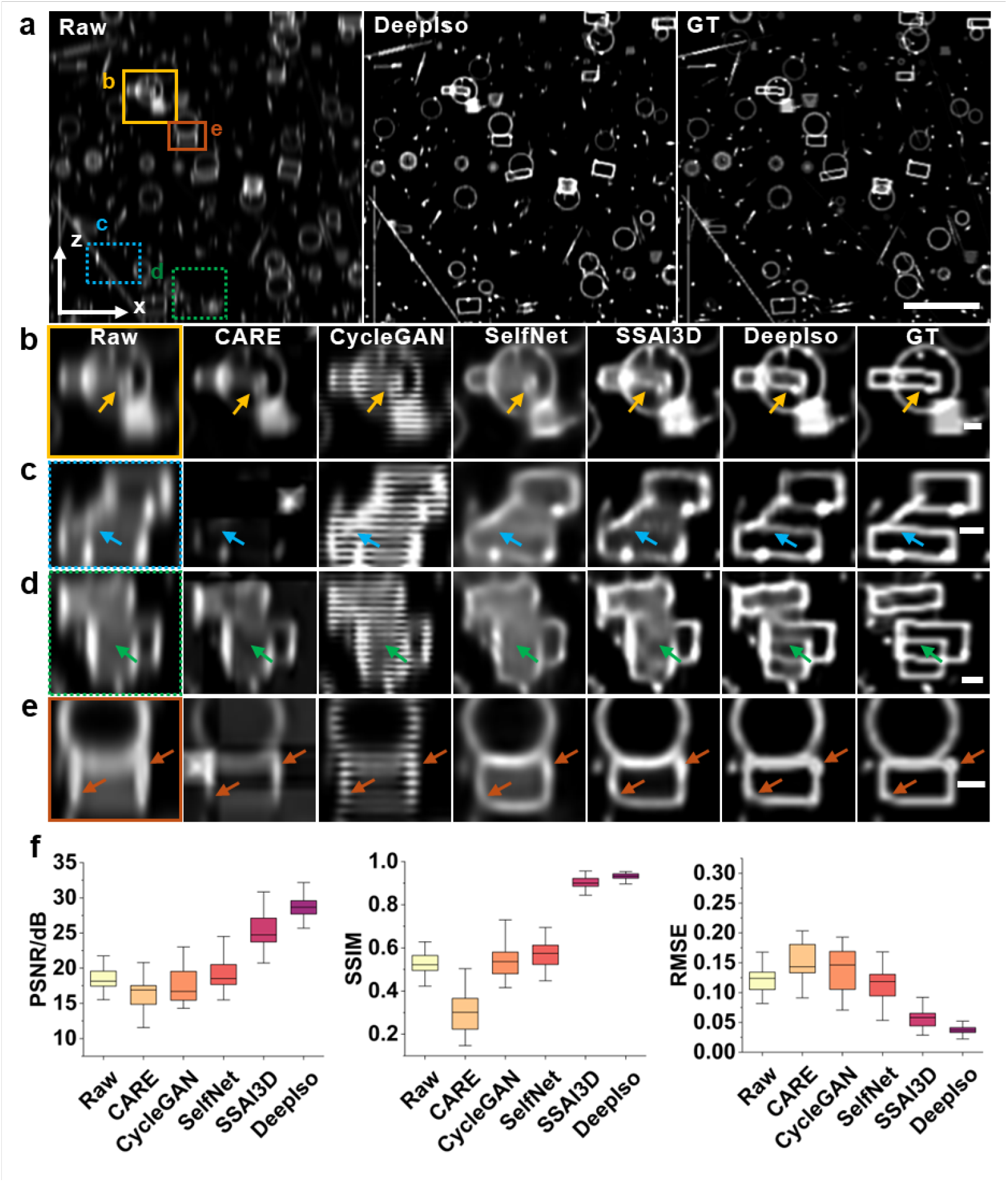
DeepIso achieves superior isotropic restoration on simulated volumetric data. **a**, Maximum-intensity projections showing that DeepIso successfully restores isotropic structure from anisotropic input, producing results that closely match the ground truth (GT). **b–e**, Magnified views of the boxed regions in **a**, comparing DeepIso with state-of-art methods, including CARE, CycleGAN, SelfNet, and SSAI3D. DeepIso effectively recovers fine structural details (arrows), preserving axial continuity while avoiding introducing artifacts or hallucinations. **f**, Quantitative evaluation using peak signal-to-noise ratio (PSNR), structural similarity index (SSIM), and root mean square error (RMSE) demonstrates that DeepIso consistently outperforms existing methods in both fidelity and isotropic accuracy. Means and standard deviations are obtained from N =31 images. Scale bars: **a** 10μm; **b-e** 1μm.

Quantitative analysis further confirmed the superior reconstruction fidelity provided by DeepIso (**Fig. 2f**). Among all evaluated methods, DeepIso achieved the highest peak signal-to-noise ratio (PSNR) and structural similarity index (SSIM), alongside the lowest root mean square error (RMSE). These metrics indicate that DeepIso produces restorations with minimal distortion and noise amplification while maintaining accurate volumetric geometry. Collectively, the results on synthetic benchmarks demonstrate that DeepIso generalizes robustly to anisotropic inputs and achieves class-leading isotropic restoration accuracy across a wide range of 3D structural motifs.

### 2.3. DeepIso performance validation with hardware-based isotropic references

To evaluate the capability of DeepIso to recovering fine axial details in experimental data, we applied it to volumetric datasets acquired by super-resolution and light-sheet modalities, where we could obtain near-isotropic ground truth (**Fig. 3**). In a first example, we used DeepIso to restore three-dimensional structured illumination microscopy (3D-SIM)^9,39^ datasets of fixed U2OS cells labeled with Alexa Fluor 488–immunolabeled microtubules. The restoration’s improved axial resolution better revealed the microtubule filament network in lateral and orthogonal views (**Fig. 3a,b**). Compared to reconstructions obtained with CARE, CycleGAN, SelfNet, and SSAI3D, DeepIso substantially improved the filament bundle continuity in axial views, closely resembling the ground truth reference provided by the hardware-based axial resolution enhancement approach, four-beam SIM (4B-SIM)^9^. Inspection of selected regions (**Fig. 3c**) confirmed that DeepIso effectively recovered submicron axial features and suppressed stripe-like or granular artifacts that persisted in other learning-based methods or hardware-based methods (**Supplementary Fig. 3**). Quantitative evaluation using normalized cross-correlation (NCC) in the axial direction (**Fig. 3d**) also demonstrated that DeepIso achieved the highest structural consistency with the ground truth. The corresponding normalized axial intensity profiles (**Fig. 3e**) also showed that DeepIso closely matched the 4B-SIM profile, indicating faithful recovery of axial modulation depth and resolution.

**Fig. 3.**
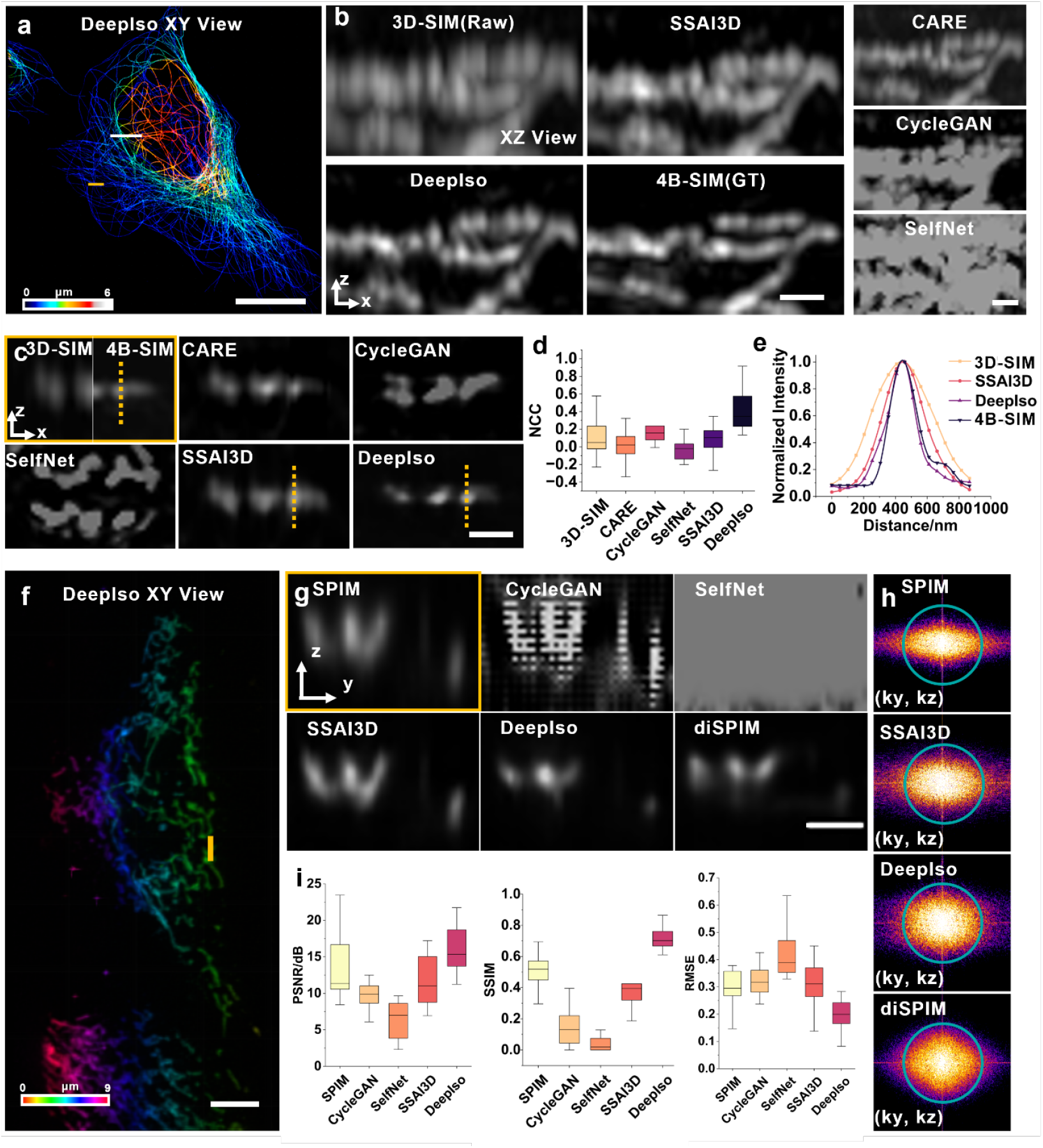
DeepIso performance benchmarking with hardware-based isotropic references. **a**, Depth-encoded rendering of fixed U2OS cells with immunolabeled microtubules restored by DeepIso (color encodes axial position, z). The raw volume was acquired by 3D-SIM. The result displays continuous filament networks with restored axial isotropy. **b**, Comparison of axial view (XZ) MIP (through a y depth range of 0.4 μm around the white line in **a**) among 3D-SIM (raw), CARE, CycleGAN, SelfNet, SSAI3D, DeepIso, and 4B-SIM (ground truth, GT). DeepIso delivers markedly improved axial resolution closely approaching that of hardware-based near-isotropic references of 4B-SIM. **c**, Representative ROI of single slice (indicated by the yellow line in **a**) illustrates the restoration by different algorithms, demonstrating DeepIso’s outperformance over other methods. **d**, Normalized axial intensity profiles (corresponding to the yellow dashed lines in **c**) showing that DeepIso best matches the 4B-SIM ground truth. Note the SelfNet and CycleGAN profiles are not necessarily displayed due to their overextended widths. **e**, Normalized cross-correlation (NCC) to the 4B-SIM ground truth quantifying axial resolution enhancement across all compared methods. Means and standard deviations are obtained from N = 26 ROIs. **f**, DeepIso restoration of Live U2OS cells transfected with mEmerald-Tomm20 from a single view raw image acquired with dual-view light sheet microscopy. **g**, Representative axial view (YZ) slice (indicated by the yellow line in **f**) comparing SPIM (single view, raw), CycleGAN, SelfNet, SSAI3D, DeepIso, and diSPIM (ground truth reference, the fusion of physical dual-view acquisitions). DeepIso substantially enhances axial resolution under high-NA illumination, approaching hardware-based multiview light sheet (diSPIM) performance. **h**, Fourier spectra of SPIM, SSAI3D, DeepIso, and diSPIM images, demonstrating the expanded axial-frequency support recovered by DeepIso. The blue circles indicate 1/330 nm^-1^ spatial frequency. **i**, Quantitative comparison of PSNR, SSIM, and RMSE across all methods, confirming that DeepIso consistently achieves the highest reconstruction fidelity. Means and standard deviations are obtained from N = 18 ROIs. Scale bars: **a** 16 μm; **b** 0.5 μm; **c** 0.5 μm; **f** 8 μm; **g** 2 μm.

Next, we tested DeepIso on mitochondria in live U2OS cells transfected with mEmerald–Tomm20, acquired by dual-view light sheet microscopy^12,13,22^, which offers near-isotropic resolution by jointly deconvolving the raw data from two orthogonal views (**Fig. 3f**). We evaluated the performance of DeepIso on single-view stacks acquired at 0.8 detection NA (SPIM), against which we also had ground truth dual-view reconstructions (diSPIM, ground truth). Even under this relatively modest NA, DeepIso produced isotropic restorations with better mitochondrial membrane continuity and network morphology compared to the input data. In representative axial slices (**Fig. 3g**), DeepIso outperformed CycleGAN, SelfNet, and SSAI3D, recovering fine axial mitochondrial details that remained blurred, fragmented, or artifact-prone in other methods, and more closely approached the dual-view reconstruction reference. Fourier-domain analysis (**Fig. 3h**) further confirmed the greater axial-frequency support resembling diSPIM, consistent with improved axial resolution and contrast. Quantitatively, DeepIso achieved the highest PSNR and SSIM and the lowest RMSE across all tested methods (**Fig. 3i**), indicating superior restoration fidelity and structural isotropy relative to the diSPIM reference. Together, these results demonstrate that DeepIso robustly enhances axial resolution in both 3D-SIM of immunolabeled U2OS microtubules and single-view SPIM data of live U2OS mitochondria, narrowing the gap between anisotropic raw inputs and isotropic ground-truth references.

### 2.4. DeepIso achieves isotropic resolution across multi-modality imaging

We next examined whether DeepIso could generalize to conventional diffraction-limited imaging modalities, focusing on confocal fluorescence microscopy, which typically suffers from pronounced axial–lateral resolution anisotropy. We evaluated DeepIso on a volumetric Thy1-eYFP mouse neuron dataset^33^ acquired by confocal fluorescence microscopy with optical sectioning^33^. As reported for this dataset, optical sectioning produces a stark contrast between lateral and axial resolution, with highly anisotropic voxel sampling (0.207×0.207×1 μm^3^) in the raw volume. DeepIso resampled the volume and produced near isotropic resolution with isotropic voxels (0.207×0.207×0.207 μm^3^) (**Fig. 4a**). The maximum-intensity projections of axial views (**Fig. 4b,d**) revealed markedly improved axial continuity and contrast compared with the original confocal inputs. Magnified subregions and corresponding Fourier spectra (**Fig. 4c,e,f**) further indicate enhanced recovery of axial high-frequency components and expanded axial-frequency support in the Fourier domain, enabling finer delineation of densely interwoven neuronal processes with consistent morphology across viewing directions. Line profiles extracted along the axial direction (**Fig. 4g,h**) showed sharper peak separation and reduced axial broadening after DeepIso restoration, while local region analysis (**Fig. 4i**) corroborated isotropic frequency recovery across the dataset.

**Fig. 4.**
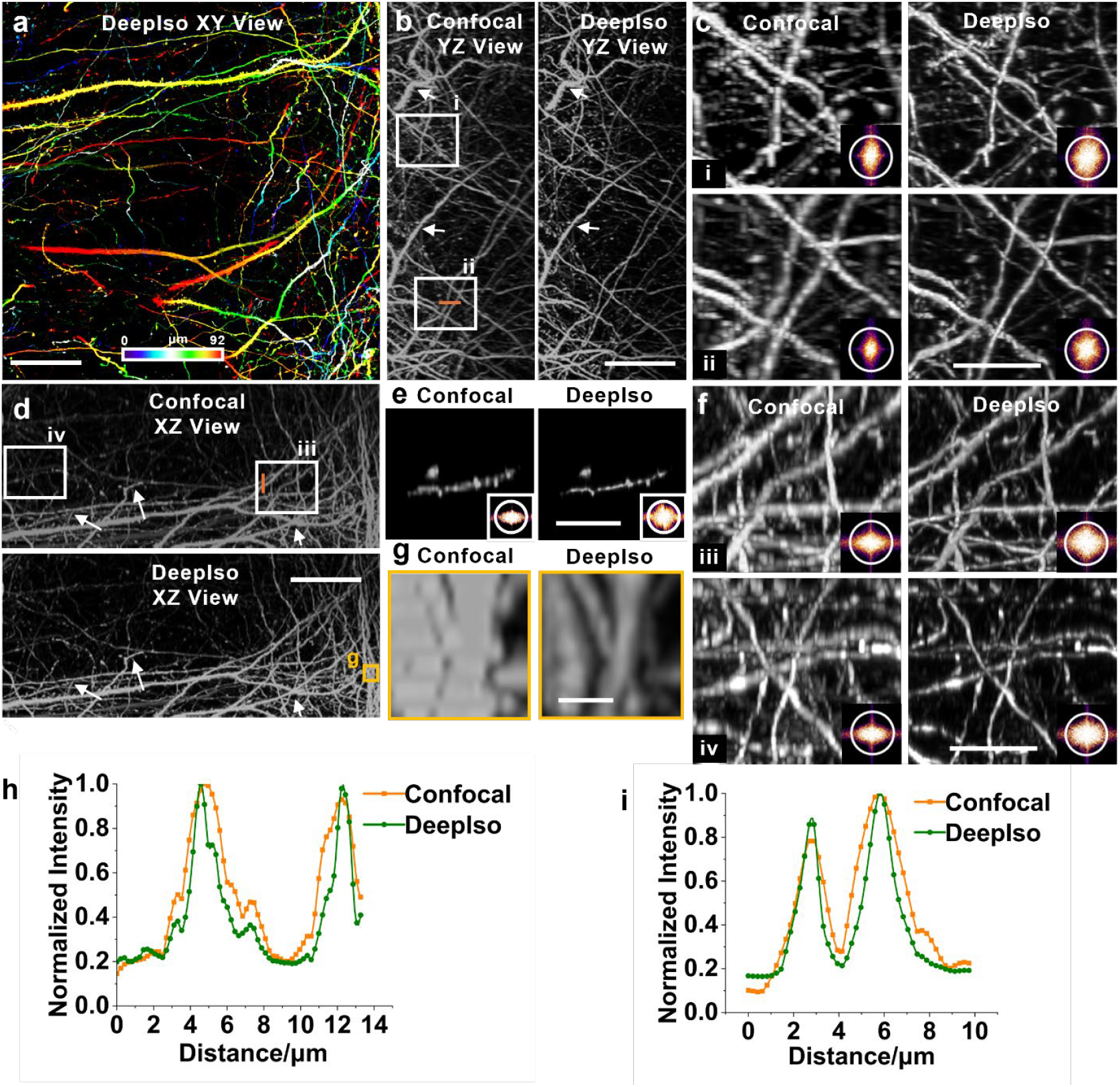
DeepIso enhances isotropic resolution in dense, highly entangled neuron samples acquired with confocal microscopy. **a**, Depth-encoded rendering of mouse Thy1 neurons acquired by confocal microscopy and restored by DeepIso, showing dense and continuous neuronal distribution in lateral view (XY). **b**, Axial view (YZ) maximum-intensity projections (MIP) from confocal and DeepIso reconstructions, revealing enhanced axial continuity and finer filament recovery. **c**, Magnified ROIs from **b** and corresponding Fourier spectra, indicating extended axial frequency support after DeepIso restoration. The white circles indicate 1/640 nm^-1^ spatial frequency. **d**, Another axial view (XZ) MIPs highlighting improved axial resolution across neuronal networks. **e**, Representative axial slice (XZ at y=147.8 μm) showing local ROI and its corresponding Fourier spectra, further demonstrating restoration of high-frequency axial information. **f**, Magnified ROIs from **d** and their Fourier spectra confirming isotropic resolution gain. **g**, Small-area axial view (XZ) MIPs illustrating consistent recovery of axial resolution. **h–i**, Normalized intensity profiles along selected YZ and XZ lines (orange lines) from **b** and **d**, respectively, showing sharper peak separation and reduced axial broadening after DeepIso processing. Scale bars: **a** 20 μm; **b** 20 μm; **c** 10 μm; **d** 20 μm; **e** 10 μm; **f** 10 μm; **g** 2 μm.

We further evaluated DeepIso on experimental super-resolution 3D-SIM datasets^9^. In bacterial membrane imaging (**Fig. 5a–d**), DeepIso markedly improved axial contrast and reduced axial elongation, enabling clearer delineation of the two closely spaced membrane layers within individual bacteria, which appear axially smeared in the native 3D-SIM volume. Consistent with the visual restoration, Fourier analysis (**Fig. 5b**) showed an expanded axial-frequency support after DeepIso, and the corresponding magnified view comparisons and line profiles (**Fig. 5c,d**) confirmed enhanced peak separation and reduced axial broadening.

**Fig. 5.**
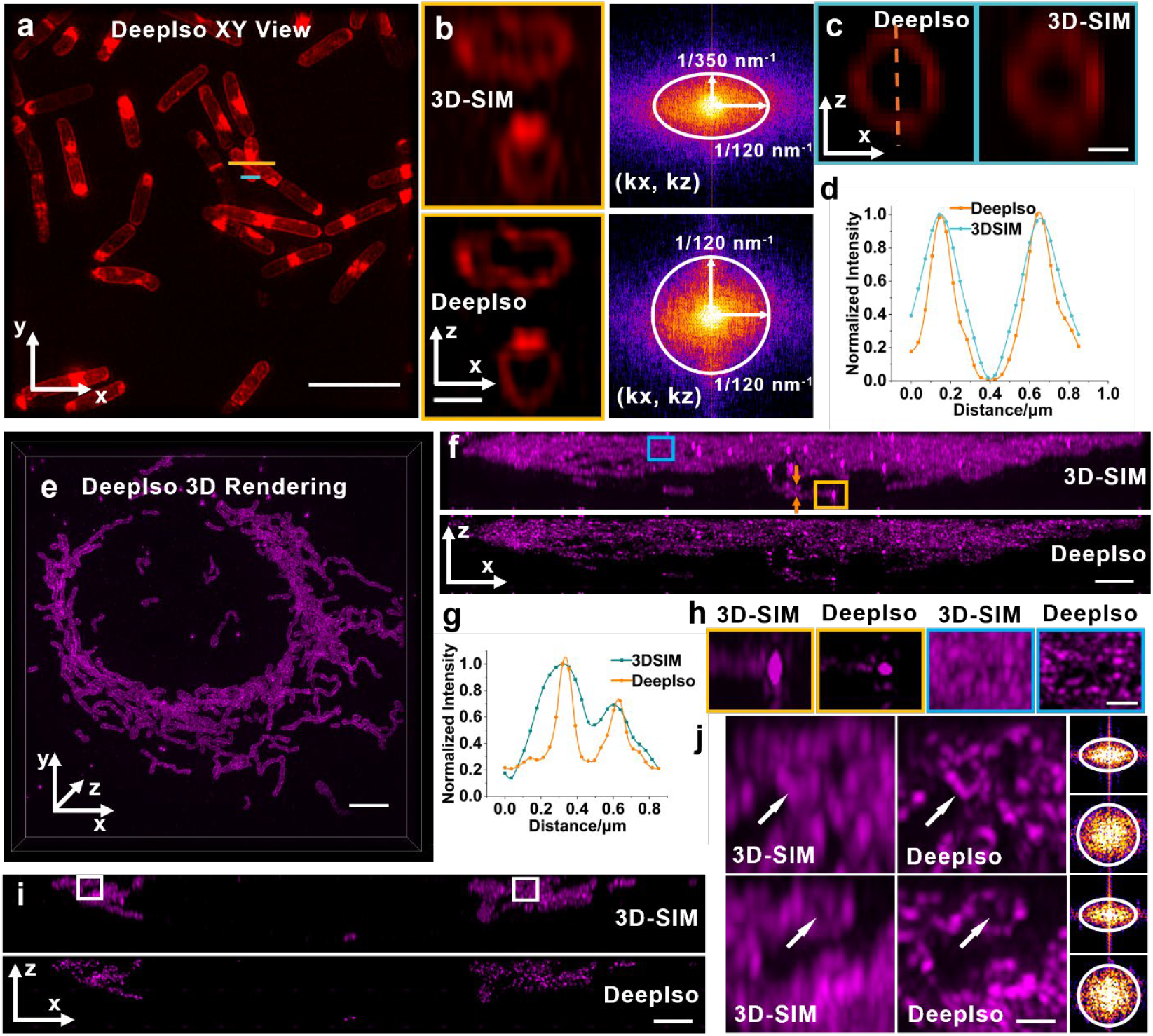
DeepIso improves isotropy and preserves fine structures in 3D SIM imaging. **a**, Lateral view (XY) maximum-intensity projection (MIP) of bacterial membranes restored by DeepIso, showing improved continuity and isotropic membrane morphology. **b**, Representative ROI of axial view (XZ) slice (indicated by the yellow line in **a**) and corresponding Fourier spectra illustrating extended axial-frequency support after DeepIso restoration compared with 3D-SIM. **c**, Another ROI of lateral slice (indicated by the cyan line in **a**) showing clearer axial separation of membrane contour by DeepIso. **d**, Normalized intensity profiles along the dashed line in **c**, demonstrating enhanced axial contrast and sharper separation of features. **e**, 3D rendering of U2OS outer mitochondrial membranes restored by DeepIso showing continuous network morphology. **f**, Axial view (XZ) MIPs of 3D-SIM raw data and DeepIso restorations demonstrating superior axial resolution recovery by DeepIso. **g**, Normalized intensity profiles along the region marked in **f**, confirming axial improvement. **h**, Magnified ROIs from **f** showing detailed membrane substructures. **i**, Lateral view (XZ) MIP (through y depth between 19.223 μm and 21.268 μm) of 3D-SIM raw and DeepIso restoration, revealing consistent isotropic recovery across the volume. **j**, Representative ROIs from **i** and corresponding Fourier spectra showing restoration of high-frequency axial components (arrows) by DeepIso. The white circle indicates the lateral cutoff frequency (1/330 nm^-1^). Scale bars, **a** 4 μm; **b** 0.5 μm; **c** 0.25 μm; **e** 4 μm; **f** 2 μm; **h** 0.2 μm; **i** 2 μm; **j** 0.2 μm.

We next applied DeepIso to 3D-SIM images of U2OS outer mitochondrial membranes^9^ (**Fig. 5e–j**). DeepIso restored a continuous and morphologically coherent mitochondrial network in lateral views (**Fig. 5e**) and substantially sharpened mitochondrial boundaries along the axial direction (**Fig. 5f,i**). Line profiles (**Fig. 5g**) revealed increased axial modulation depth and improved separation of adjacent structures, while enlarged ROIs (**Fig. 5h,j**) highlighted recovery of fine axial details with reduced striping artifacts. Fourier spectra from selected regions (**Fig. 5j**) further supported the restoration of higher axial-frequency components compared with conventional 3D-SIM.

Together, these results indicate that DeepIso can enhance axial resolution and 3D structural fidelity across both diffraction-limited and super-resolution modalities without requiring explicit PSF specification or manual modality-specific parameter tuning, while using the same training and adapting protocol per modality.

### 2.5. DeepIso facilitates reliable segmentation and cell tracking for time-lapse data

To further demonstrate the applicability of DeepIso to long-term volumetric imaging, we applied it to dual-view light sheet microscopy datasets of zebrafish embryos expressing Lyn–eGFP under the claudin B (cldnb) promoter^22^, which labels epithelial cell membranes around the lateral-line primordium (**Fig. 6a**). The single view raw data (SPIM) exhibited characteristic axial blurring and discontinuities in membrane junctions, particularly along the apical–basal direction. DeepIso restoration significantly improved axial continuity and junctional delineation, producing volumetric restoration comparable to the two-view reconstruction results (diSPIM), but without requiring explicit two-view data acquisition and fusion (**Fig. 6b**).

**Fig. 6.**
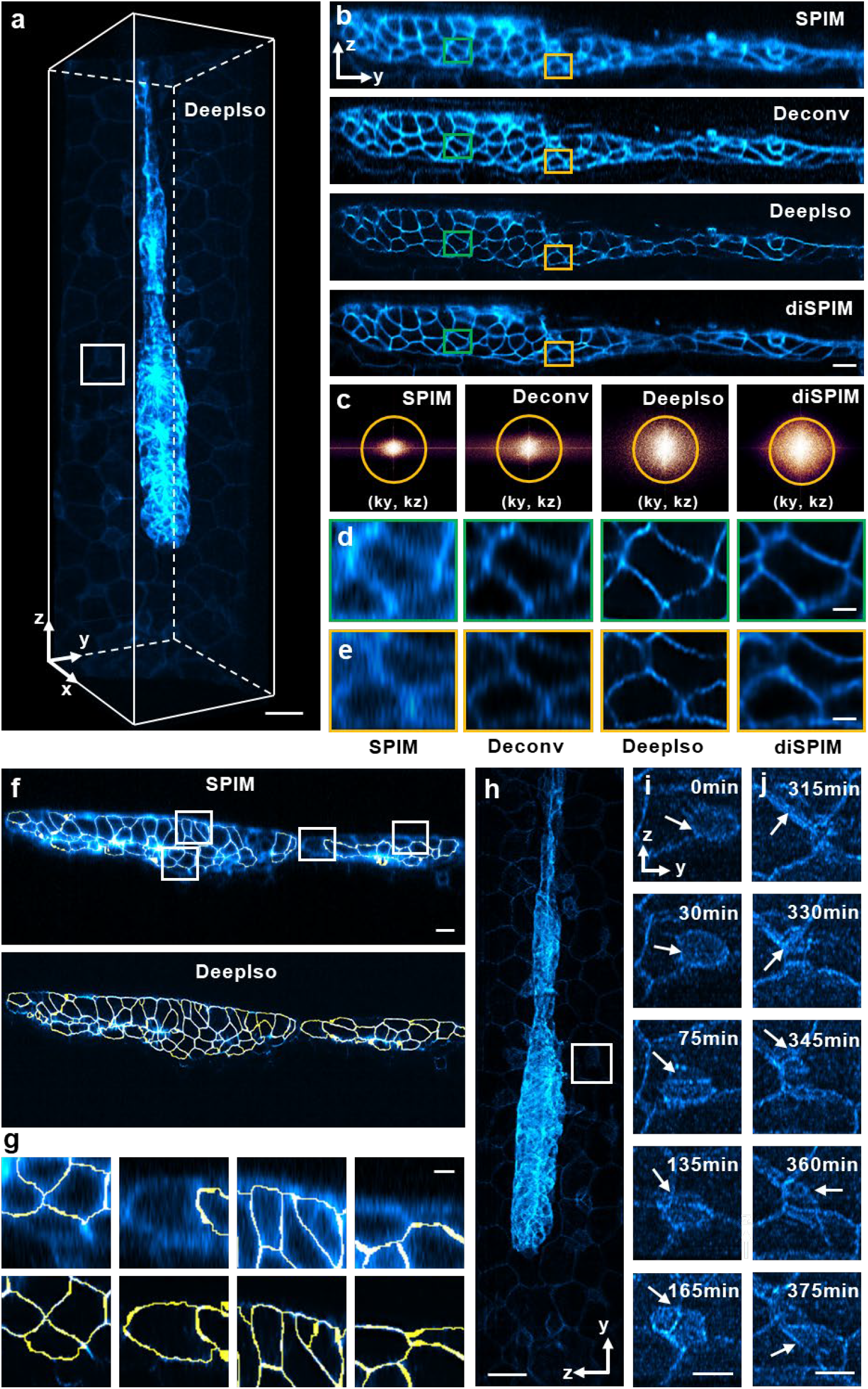
DeepIso enables reliable segmentation and cell tracking for time-lapse data by improving structure fidelity. **a**, Three-dimensional rendering of a 32-h post-fertilization zebrafish embryo expressing Lyn–eGFP under the claudin B (cldnb) promoter, highlighting cell membranes within and surrounding the lateral-line primordium. A single view raw volume from light sheet microscopy was restored with DeepIso. **b**, Representative axial slices (YZ at x=62.56 μm) from single-view raw (SPIM), single view deconvolution (Deconv), DeepIso restoration, and the fusion result of two-view acquisitions (diSPIM, ground truth), showing markedly improved axial resolution and junctional details in DeepIso. **c**, Corresponding Fourier spectra for different methods. The yellow circles indicate 1/330 nm^-1^ spatial frequency. The extended axial-frequency support recovered by DeepIso outperforms conventional SPIM and deconvolution, approaching the isotropic resolution of diSPIM, which fuses two-view acquisitions. **d, e**, Magnified ROIs from **b** showing isotropic restoration of epithelial boundaries and enhanced contrast. **f**, Segmentation of primordium (YZ at x=55.7 μm) illustrating DeepIso restoration provides more accurate epithelial boundary delineation than SPIM raw slice. **g**, Representative ROIs from **f** highlighting continuous and morphologically preserved membranes by DeepIso (bottom) over raw (top). **h**, Time-lapse axial view (YZ) MIP reconstructed by DeepIso, visualizing apical and basal surfaces of the primordium with high axial fidelity (time point 30 min is displayed). **i, j**, Two independent ROIs tracking distinct immune cells (arrows) migrating between surrounding skin cells from 0 – 375 min by DeepIso. Scale bars, **a** 20 μm; **b** 10 μm; **d** 2 μm; **e** 2 μm; **f** 10 μm; **g** 10 μm; **h** 20 μm; **i** 8 μm; **j** 8 μm.

Fourier-domain analysis (**Fig. 6c**) revealed that DeepIso expanded the recoverable axial-frequency range relative to the single view raw (SPIM) and corresponding deconvolution (Deconv) images, reflecting enhanced resolution along the optical axis. Enlarged ROIs (**Fig. 6d,e**) demonstrated smooth and isotropic recovery of epithelial boundaries with enhanced contrast, allowing clear visualization of individual cell membranes within the densely packed epithelial layers. Segmentation of the DeepIso restored volumes (**Fig. 6f,g**) further confirmed accurate and continuous delineation of cell boundaries across the primordium, in contrast to the fragmented or partially fused junctions in the original single view data.

Importantly, DeepIso facilitated manual tracking and automatic segmentation in long-term time-lapse imaging. In the 6.25-hour light sheet microscopy recording, DeepIso restored isotropic morphology and continuity of the migrating primordium (**Fig. 6h**) while preserving dynamic subcellular features. Two distinct regions of interest (**Fig. 6i,j**) revealed two immune cells (arrows) migrating between surrounding skin cells: one cell in the period of 0–165 minutes (**Fig. 6i**) and another in the period of 315–375 minutes (**Fig. 6j**), highlighting DeepIso’s ability to maintain clear cell outlines and continuous elongated structures across frames.

Together, these results demonstrate that DeepIso not only restores isotropic spatial resolution in light-sheet microscopy but also enables long-term volumetric imaging with improved spatiotemporal resolution. By recovering high-frequency axial information directly from single-view raw data, DeepIso eliminates the dependence on multi-view data acquisition and complex multiview fusion procedures, thus providing a scalable framework for high-fidelity live-cell and developmental imaging.

### 3. Discussion and conclusion

In this study, we introduced DeepIso, a self-supervised framework for restoring isotropic resolution in three-dimensional microscopy. By leveraging a two-stage training paradigm that integrates synthetic pretraining with adaptive inference, DeepIso learns modality-independent isotropic features directly from the data, eliminating the need for paired high-resolution supervision or explicit multi-view fusion. Across diverse imaging modalities, including 3D-SIM, SPIM, diSPIM, and confocal microscopy, DeepIso consistently enhanced axial resolution, improved structural isotropy, and preserved fine morphological details, while minimizing artifacts and noise amplification.

Unlike previous learning-based restoration approaches that rely on explicit supervision (e.g., CARE) or adversarial domain translation (e.g., CycleGAN, SelfNet), DeepIso performs internal adaptation through latent-space consistency. The encoder–decoder pair (IsoEnc/IsoDec) remains fixed during adaptation, ensuring stable feature representation and preventing overfitting to dataset-specific noise patterns. This design enables robust generalization across different imaging systems, as evidenced by DeepIso’s strong performance on both simulated benchmarks and experimental datasets, including live-cell and organismal imaging.

A key advantage of DeepIso lies in its ability to recover high-frequency axial components from single-view anisotropic inputs, without the need for external calibration data. Fourier-domain analysis confirmed that DeepIso effectively expands the axial-frequency support, thereby restoring isotropic point-spread responses and volumetric contrast. This improvement in isotropy directly benefits downstream analyses, such as segmentation and morphological quantification. Beyond static imaging, DeepIso demonstrated substantial utility in time-lapse volumetric analysis. In diSPIM recordings of developing zebrafish embryos, the method restored continuous epithelial boundaries and facilitates manual tracking of immune-cell migration over extended durations with high spatiotemporal fidelity (**Fig. 6**). Such capability underscores DeepIso’s potential for studying dynamic biological processes that require both gentle illumination and precise 3D reconstruction, such as tissue morphogenesis, organ development, and intracellular transport. This application highlights DeepIso’s robustness in both high-resolution anatomical imaging and time-resolved dynamic analysis.

Although DeepIso demonstrates strong isotropic restoration performance under matched imaging conditions, several limitations remain. First, its generalization across heterogeneous biological samples is inherently constrained. In practice, different specimen types exhibit substantially different structural organizations, signal-to-noise characteristics, and anisotropic degradation patterns. Consequently, optimal performance typically requires dataset-specific training or adaptation, rather than reliance on a single universal model. This limitation reflects a broader challenge in microscopy image restoration, where strong domain shifts arise from biological diversity and optical variability. Future work may explore generalized pretrained representations^40,41^ or meta-learning strategies^42^ to improve cross-sample generalization with reduced retraining cost. Second, DeepIso introduces additional computational cost associated with its multi-component training framework. For example, in our experiments on confocal Mouse Thy1 neuron data, training of the IsoNet module required approximately 4.8 hours, while training of the IsoEnc/IsoDec module required approximately 6.0 hours on a single NVIDIA RTX 4080 SUPER GPU. The adapting-inference stage converged substantially faster, requiring approximately 13 minutes for 12,000 iterations. Notably, the training of IsoNet and IsoEnc/IsoDec is structurally independent. When computational resources permit, these two components can be trained concurrently on separate GPUs, which can effectively reduce the overall wall-clock training time.

Once training and adaptation are complete, DeepIso can be directly applied to raw biological image volumes in a purely feed-forward manner, analogous to conventional end-to-end restoration networks such as CARE^23^ or RCAN3D^26^. In this inference stage, no additional optimization or parameter updating is required, and large volumetric datasets can be processed efficiently using standard block-wise reconstruction strategies. Third, DeepIso conceptually could be extended to laterally anisotropic data, but this would require redesigning the network to recover isotropy from only one high-resolution lateral direction. We expect substantial additional development and validation, so we leave this as future work and only note it here. Finally, future extensions of DeepIso may benefit from integration with multi-view fusion strategies or physics-informed priors, particularly for complex specimens exhibiting strong light scattering, refractive index heterogeneity, or severe optical aberrations. Such hybrid designs may further enhance reconstruction robustness and extend applicability to challenging biological imaging scenarios.

Overall, DeepIso provides a unified and scalable framework for isotropic 3D reconstruction across a broad range of fluorescence microscopy modalities. Its self-supervised learning scheme, internal adaptation, and strong cross-domain generalization make it a powerful tool for next-generation volumetric imaging, which can bridge the gap between diffraction-limited acquisitions and isotropic, artifact-free reconstructions suitable for quantitative biological analysis. DeepIso thus offers a significant step forward in enabling high-quality, high-fidelity imaging in both static and dynamic biological studies.

## Methods

The DeepIso framework adopts a modular architecture composed of an isotropic restoration backbone (IsoNet) and a latent feature encoder–decoder module (IsoEnc/IsoDec). The model is first pretrained on synthetically generated anisotropic–isotropic patch pairs to establish a stable isotropy-aware mapping. During adapting inference, the latent feature learned by IsoEnc is combined with experimental observations through IsoDec, enabling unsupervised refinement under real imaging conditions. This two-stage strategy ensures both reconstruction fidelity and robustness to system-dependent anisotropy.

### Network architecture

IsoNet was implemented as a compact residual channel attention network (RCAN) backbone^**26**^. A single channel input was linearly standardized from 0 to 1 into minus 1 to 1, passed through a 3 by 3 feature extraction layer with 32 channels, and processed by five residual groups, each containing five residual channel attention blocks with reduction ratio 8. Each block used two 3 by 3 convolutions with a ReLU after the first, followed by channel attention based on global average pooling and 1**×**1 bottleneck layers. A long skip connection bridged the residual group stack, and a final 3**×**3 convolution produced a single channel output that was mapped back to 0 to 1.

IsoEnc and IsoDec were implemented jointly as a two-branch RCAN based architecture. Both inputs were first linearly standardized from 0 to 1 into minus 1 to 1. IsoEnc encoded the first input using an RCAN style encoder with 64 feature channels and three residual groups, and produced a 1024 dimensional latent representation via global average pooling followed by a 1 by 1 convolution. IsoDec conditioned the reconstruction on both the latent code and the second input. The latent code was expanded by a 1 by 1 convolution and bilinear upsampling to match the spatial patch size, then concatenated with features extracted from the second input by a symmetric RCAN stem. The fused features were refined by three residual groups and projected to output, which was mapped back to 0 to 1 by the inverse linear transform.

#### 1) IsoNet fomulation

Let the input patch be ***x*** ∈ [**0**,**1**]^***H***×***W***^, a linear standardization is first applied

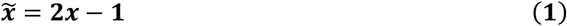

Let 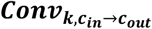 (·) denote as 2D convolution with kernel size ***k*** × ***k, ReLU*** (·) denote ReLU, and *σ*(·) denote sigmoid. For a feature map **u** ∈ ℝ^***C***×***H***×***W***^, global average pooling is

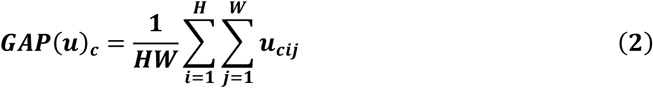

##### Channel attention

With reduction ratio ***r***, the channel attention weights are computed as

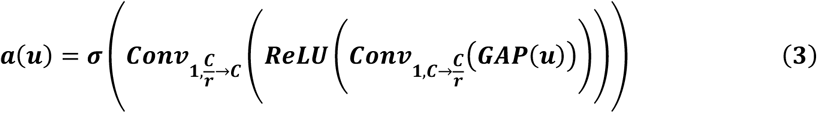

and applied by channel wise reweighting

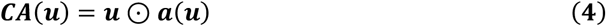

where ⊙ denotes elementwise multiplication with broadcasting over spatial dimensions.

##### Residual channel attention block

With residual scaling α, one residual channel attention block is

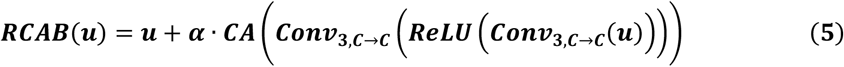

##### Residual group

A residual group with ***B*** blocks is defined as

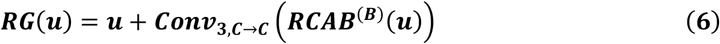

where ***RCAB***^(***B***)^ denotes a sequential composition of ***B*** RCAB modules.

##### RCAN mapping

Let the initial feature extraction be

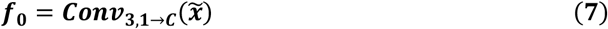

After stacking ***G*** residual groups,

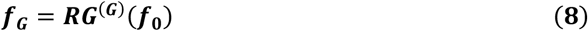

the long skip connection and reconstruction head are

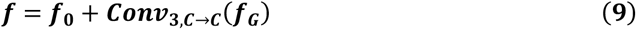

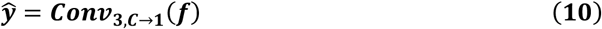

followed by inverse scaling to the range [0, 1]

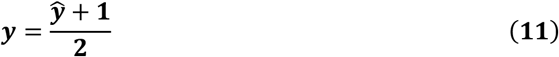

In our IsoNet configuration, ***C***=32, ***B***=5, ***G***=5, ***r***=8, and α=1.0.

#### 2) IsoEnc and IsoDec fomulation

The IsoEnc and IsoDec models take two inputs ***x, y*** ∈ [**0, *1***]^***H***×***W***^ and applies the same linear standardization

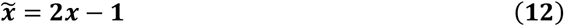

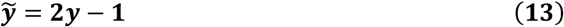

##### IsoEnc

Let ***ψ***(·) denote the encoder stem implemented by reflection padding, a 7×7 convolution, and a LeakyReLU. The encoder feature map is

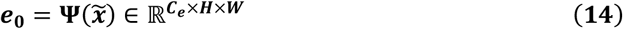

Let 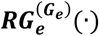 denote the stack of encoder residual groups, each built from residual channel attention blocks with a local residual connection. The encoder output before pooling is

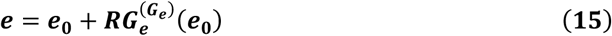

latent features are then produced by global average pooling followed by 1×1 projections to latent dimension ***D***

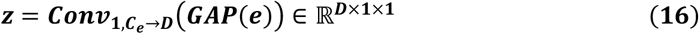

##### IsoDec

Let ***UP***(·) denote bilinear upsampling to size ***H*** × ***W***. The latent features are expanded to a spatial feature map

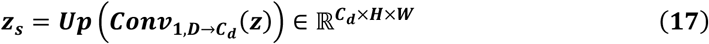

Let **Φ**(·) denote the decoder conditioning stem applied to 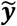, implemented by reflection padding, a 7×7 convolution, and a LeakyReLU

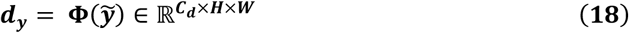

The two feature maps are concatenated along channels are refined by a stack of decoder residual groups 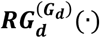

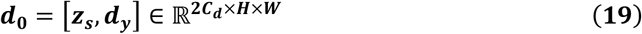

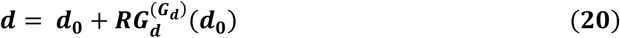

A final reconstruction head ***Ω***(·), implemented by refection padding and a 7×7 convolution, produces the output

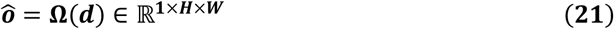

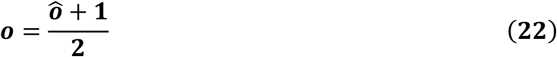

##### Overall mapping

The complete model is

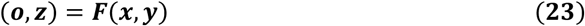

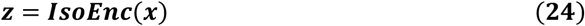

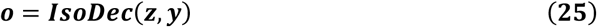

In our configuration, ***C***_***e***_= ***C***_*d*_=64, ***G***_***e***_=***G***_*d*_=3, and ***D***=1024.

### Training losses

During pretraining, pixelwise supervision was imposed using the Charbonnier loss, a differentiable and robust variant of the L1 norm. For a predicted image ***I***_***pred***_ and its reference ***I***_***ref***_, the loss was computed as

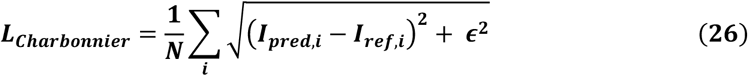

where ***i*** indexes pixels, ***N*** denotes the total number of pixels, and *ϵ* is a small constant for numerical stability. The loss was averaged over all pixels using mean reduction and applied with a weight of 1.0.

Pixelwise supervision during adapting inference was implemented with a mean squared error criterion. Given a network prediction ***I***_***pred***_ and the corresponding reference ***I***_***ref***_, the objective was evaluated as

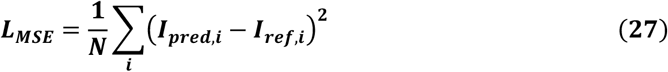

in which ***i*** denotes the pixel index and ***N*** is the total number of pixels. Mean reduction was used, and the resulting loss term was assigned a weight of 1.0.

### Training details

The DeepIso framework was implemented in PyTorch and optimized using adaptive moment estimation (Adam) with ***β1*** =0.9 and ***β2*** =0.99. Training was performed for 100,000 iterations in the pretraining stage and 12,000 iterations in the adapting-inference stage. An exponential moving average (EMA) of the model parameters with a decay rate of 0.999 was maintained to promote stable convergence. The initial learning rate started at **1** × **10**^−**4**^ and scheduled using cosine annealing over the full training course, gradually decaying to a near-zero minimum toward the end of training. All experiments were conducted on a workstation equipped with Nvidia GeForce RTX 4080 SUPER GPU (with 16 GB of RAM).

In the pretraining stage, we set the batch size to 8, initial learning rate to **1** × **10**^−**4**^ and the worker number per GPU to 6. During the adapting inference stage, we set the batch size to 8, initial learning rate to **5** × **10**^−**7**^ and the worker number per GPU to 6.

### INFERENCE and block-wise reconstruction strategy

To accommodate large volumetric datasets exceeding GPU memory limits, DeepIso inference was performed in a block-wise manner with controlled spatial overlap between adjacent tiles. Each 3D input volume was first divided into overlapping sub-volumes (typically 64^3^–128^3^ voxels per block) with an overlap margin of 16–32 voxels along each axis. The overlapping regions were smoothly blended using a linear weighting window to avoid boundary discontinuities after reconstruction.

Before inference, anisotropic raw data were aligned through the same pixel-alignment procedure used during training, i.e., one spatial dimension was downsampled to match the axial pixel size and subsequently upsampled to the lateral sampling interval. The aligned volumes were then processed by the pretrained IsoNet adapted through Adapting Inference Stage.

All convolutional layers operated on isotropic receptive fields, ensuring that the network preserved structural consistency across the axial and lateral dimensions. The encoder and decoder components (IsoEnc/IsoDec) remained frozen during inference, maintaining a stable latent-space mapping learned during pretraining. The intermediate IsoNet weights, fine-tuned during the adaptive stage, handled residual anisotropy and dataset-specific characteristics such as refractive mismatch, optical aberration, and SNR variation.

The restored sub-volumes were finally stitched into the full 3D isotropic reconstruction using weighted averaging in the overlap regions, producing seamless volumes suitable for quantitative downstream analyses such as segmentation, tracking, and Fourier-domain isotropy evaluation.

### Dataset preparation for pretraining

In Stage-1: Pretraining, a paired pretraining dataset was constructed by extracting 2D patches from 3D TIFF volumes and generating two corresponding representations for each patch: (i) a laterally well-resolved reference patch, which serves as a proxy for isotropic lateral resolution, and (ii) an anisotropic patch, synthetically degraded to emulate direction-dependent resolution loss. In practice, the reference patches were sampled from lateral view and treated as the high-quality target, whereas the anisotropic counterparts were produced through controlled blurring and optional resampling so that the paired data explicitly encode an isotropic-to-anisotropic mapping.

All volumes were loaded from the input directory and, unless otherwise stated, intensity-normalized by percentile-based^23^ linear rescaling to map values into [0, 1], with optional clipping to suppress outliers. Patch extraction was performed slice-wise along the axial dimension. For each *z*-slice, a fixed number of square crops (default 64; 10 patches per slice) were drawn at random. Because raw microscopy volumes frequently contain extensive low-signal background, patch selection was constrained by a rejection criterion to enrich informative structures: candidate crops were repeatedly proposed (up to 1000 attempts) and retained only when their mean intensity exceeded 70% of the corresponding volume-wide mean.

The anisotropic counterparts were generated from the same reference crops using an orientation-dependent blur model, followed by an optional downsample–upsample step that mimics under-sampling. Gaussian blur kernel collection was assembled in two stages. First, a set of base Gaussian kernels with fixed support (size 31 pixels) was created by varying the standard deviation (sampled from 3 to 101 in steps of 2, retaining the first K kernels where K=*kernel_number*), and each kernel was normalized to ensure consistent scaling. Second, to reproduce directionally biased degradation, each base kernel was rotated in-plane to multiple orientations, with angles uniformly distributed over [−90^*°*^, 90^*°*^]according to *angle_number* bins; the complete kernel library was saved to disk to facilitate reproducibility. For each reference patch, the corresponding anisotropic patch was synthesized by FFT-based convolution with a selected kernel while preserving the patch size via padding. When enabled, a pixel size alignment procedure was applied by first resizing the convolved patch to a lower resolution determined by *downsampling_rate* and then interpolating it back to the original patch size, thereby introducing scale-dependent smoothing consistent with anisotropic sampling.

A deterministic patch-level split was used to obtain stable training, validation, and test partitions. Within each group of ten sampled patches, one patch was assigned to the validation set and one to the test set, while the remaining eight were used for training (approximately 8:1:1). All paired patches were saved as TIFF files into the corresponding train/val/test subdirectories.

### Dataset preparation for adapting inference

In the State-2: Adapting Inference, an unaligned patch dataset was derived directly from experimental 3D volumes so that the model could be internally adapted to real acquisition statistics without presuming cross-view correspondence. Each TIFF volume was loaded from disk and, unless stated otherwise, intensity-normalized by percentile-based linear rescaling. Values were mapped to the range from 0 to 1, and clipping was applied to suppress out-of-range fluctuations. This normalization was applied consistently across volumes to reduce scale variability while retaining specimen-specific contrast relationships.

Patch extraction was performed from orthogonal section families, since anisotropic sampling induces plane-dependent image statistics, particularly for sections that intersect the axial direction. Two sets of 2D crops were therefore generated. First, xz sections were formed by fixing the y index. Within each section, square patches of fixed size, by default 64 by 64 pixels, were randomly cropped in the xz plane, with multiple realizations drawn per section, by default 10 patches. In parallel, xy sections were formed by fixing the z index, followed by random cropping in the xy plane using the same patch size and sampling multiplicity. Because crops from the two section families were sampled independently with different random spatial coordinates, the resulting dataset was intentionally unpaired and spatially unaligned across views, which matches the adapting-inference setting in which only raw observations are available and explicit registration between orthogonal planes is not assumed.

To ensure stable data split across runs, patches were assigned to training, validation, and test sets using a deterministic indexing rule. Within each sampling loop, one out of every ten patches was routed to validation and one to test, whereas the remaining patches were used for adaptation and training, yielding an approximate 8 to 1 to 1 partition. All patches were saved as individual TIFF files into the corresponding train, val, and test directories. When available, CuPy was used to accelerate array operations; otherwise, the same protocol was executed with NumPy on CPU, without changing the sampling procedure or the resulting dataset composition.

### Evaluation metrics

Reconstruction quality was quantified using complementary error and similarity criteria computed between the predicted image ***I***_***pred***_ and its reference ***I***_***ref***_(both evaluated on the same field of view after identical cropping ande intensity scaling).

### Peak signal to noise ratio (PSNR)

PSNR was calculated from the mean squared error as

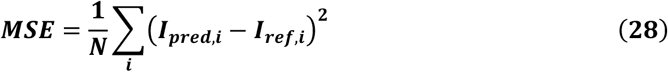

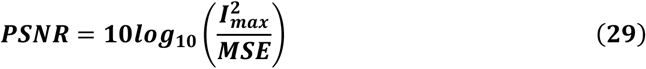

where ***i*** indexes pixels, ***N*** is the number of pixels, and ***I***_***max***_ is the maximum representable intensity (set to 1 for normalized images)

### Structural similarity index (SSIM)

SSIM was used to measure perceived structural fidelity. For two image patches ***x*** and ***y***,

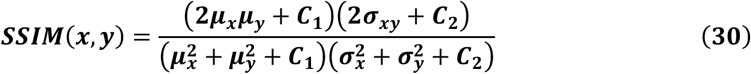

where *μ* denotes the local mean, *σ*^2^ the local variance, and *σ*_*x****y***_ the local covariance. Constants ***C***_***1***_ and ***C***_2_ were set following standard practice using the dynamic range of the images.

### Root mean squared error (RMSE)

RMSE was computed as

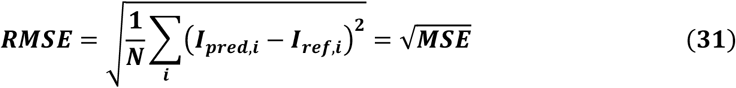

### Normalized cross correlation (NCC)

To assess global similarity up to an affine intensity shift, NCC was computed as

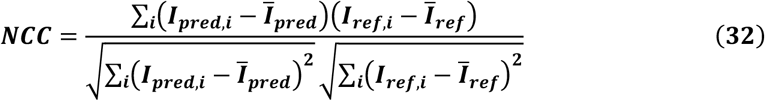

Where **Ī**_***pred***_ and **Ī**_***ref***_ are image means.

### Fourier spectrum analysis

To examine frequency-content preservation, each image was transformed using a two-dimensional discrete Fourier transform

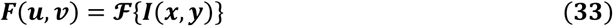

and the magnitude spectrum |***F***(***u, v***)| was visualized after shifting the zero frequency to the spectrum center using fftshift. For display, the logarithmic spectrum ***log*** (**1** + |***F***(***u, v***)|) was reported to compress the dynamic range. When required for quantitative comparison, radially averaged power spectra were computed from |***F***(***u, v***)|^2^ by binning frequencies according to their radial distance from the origin.

### Pixel size alignment

Let the anisotropic input volume be ***V*** ∈ ℝ^***H***×***W***×***z***^, with lateral pixel size **Δ**_***xy***_ and axial pixel size ***Δ***_***z***_. A pixel size alignment operator ***A***(·) was applied prior to training to equalize sampling along the selected spatial axis. Specifically, one lateral axis was first resampled to the axial spacing and then resampled back to the original lateral spacing, so that the network always observes inputs that have undergone the same effective sampling transformation.

Denote the scale factor by

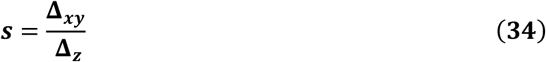

The pixel-aligned volume was obtained by a two-step interpolation

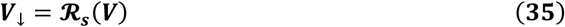

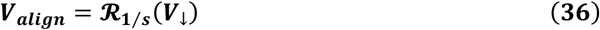

where ***R***_***s***_ denotes a one-dimensional resampling operator acting along the chosen lateral axis with scale factor ***s*** (downsampling when ***s*** < ***1*** and upsampling when ***s*** > ***1***), implemented using bicubic interpolation for 2D slices. The resulting ***V***_***align***_ preserves the original lateral grid while introducing a controlled resampling that compensates for the pixel size mismatch between axial view and lateral view, thereby enforcing uniform voxel spacing and consistent receptive-field samping across directions.

### Zebrafish segmentation

For cell segmentation in **Fig. 6f,g**, we used the MorpholibJ plugin (morphological segmentation) with the same parameter settings for SPIM and DeepIso images. Before segmentation, images were smoothed in ImageJ with a Gaussian blur (sigma = 1.5). We then applied watershed segmentation with a tolerance of 15 and a connectivity of 26.

### Competing methods

All competing methods and their quantitative results are summarized in Extended Data Tables xxx. For a fair comparison, official implementations of CARE^**23**^, CycleGAN^**32**^, SelfNet^**33**^, and SSAI3D^**34**^ were obtained from their respective GitHub repositories and retrained on the same datasets used in our experiments across imaging modalities, following the authors recommended settings whenever applicable. All evaluations were performed on a single workstation equipped with an Nvidia GeForce RTX 4080 SUPER GPU with 16 GB of memory.

## Data and code availability

The source code of this manuscript is available at https://github.com/MZ-Wei/DeepIso. The 3D-SIM and 4B-SIM datasets used in this study were obtained from the publicly available dataset released with the 4B-SIM paper^9^. The confocal microscopy dataset was obtained from the publicly available dataset reported in Ref^33^. The SPIM and diSPIM datasets were obtained from the dataset reported in Ref^22^.

## Author Contributions

H.L., M.G. and Y.L. supervised the research. M.W., P.X. and M.G. conceived the technique. M.W. and P.X. implemented DeepIso framework. M.G. and M.W. designed simulations. M.W., P.X. and Y.L. designed the validation experiment. M.W., P.X., J.L, X.F. J.Z., H.R. and R.D. performed experiments and analysis. X.L. provided 3D-SIM and 4B-SIM data and analysis. Y.H. and W.Z. provided biological insight and advice. M.W., P.X., Y.L. and M.G. wrote the manuscript with advice from all authors. All authors had access to the study.

## Acknowledgements

The work is supported by the National Natural Science Foundation of China (62427807, 62475232), the Leading Innovative and Entrepreneur Team Introduction Program of Zhejiang (2024R01001), Zhejiang Provincial Natural Science Foundation of China (LZ25F050008). We thank Hari Shroff for helpful comments on improving the manuscript.

## References

1. Stelzer, E. H. K. et al. Light sheet fluorescence microscopy. Nat. Rev. Methods Primer 1, 73 (2021).

2. Sahl, S. J., Hell, S. W. & Jakobs, S. Fluorescence nanoscopy in cell biology. Nat. Rev. Mol. Cell Biol. 18, 685–701 (2017).

3. Chen, Y., Yang, Y. & Zhang, F. Noninvasive in vivo microscopy of single neutrophils in the mouse brain via NIR-II fluorescent nanomaterials. Nat. Protoc. 19, 2386–2407 (2024).

4. Loewa, A., Feng, J. J. & Hedtrich, S. Human disease models in drug development. Nat. Rev. Bioeng. 1, 545–559 (2023).

5. Wu, Y. & Shroff, H. Faster, sharper, and deeper: structured illumination microscopy for biological imaging. Nat. Methods 15, 1011–1019 (2018).

6. Zhang, Y. et al. Super-resolution imaging of fast morphological dynamics of neurons in behaving animals. Nat. Methods 1–10 (2024) doi:10.1038/s41592-024-02535-9.

7. Fu, Y. et al. Triangle-beam interference structured illumination microscopy. Nat. Photonics https://doi.org/10.1038/s41566-025-01730-0 (2025) doi:10.1038/s41566-025-01730-0.

8. Wang, J. et al. Implementation of a 4Pi-SMS super-resolution microscope. Nat. Protoc.16, 677–727 (2021).

9. Li, X. et al. Three-dimensional structured illumination microscopy with enhanced axial resolution. Nat. Biotechnol. 41, 1307–1319 (2023).

10. Yoo, H. et al. Near-isotropic super-resolution microscopy with axial interference speckle illumination. Nat. Commun. 16, 9274 (2025).

11. He, E. et al. Boosting High-Fidelity Isotropic Super-Resolution via Image Interference Structured Illumination Microscopy with Spatial-Spectral Optimization. Laser Photonics Rev. 19, 2500178 (2025).

12. Wu, Y. et al. Spatially isotropic four-dimensional imaging with dual-view plane illumination microscopy. Nat. Biotechnol. 31, 1032–1038 (2013).

13. Kumar, A. et al. Dual-view plane illumination microscopy for rapid and spatially isotropic imaging. Nat. Protoc. 9, 2555–2573 (2014).

14. Wu, Y. et al. Simultaneous multiview capture and fusion improves spatial resolution in wide-field and light-sheet microscopy. Optica 3, 897 (2016).

15. Wu, Y. et al. Multiview confocal super-resolution microscopy. Nature 600, 279–284 (2021).

16. Wilson, T., Botcherby, E., Juškaitis, R. & Booth, M. Aberration-free refocusing in high numerical aperture microscopy. in Confocal, Multiphoton, and Nonlinear Microscopic Imaging III (2007), paper 6630_24 (Optica Publishing Group, 2007). doi:10.1364/ECBO.2007.6630_24.

17. Dean, K. M., Roudot, P., Welf, E. S., Danuser, G. & Fiolka, R. Deconvolution-free Subcellular Imaging with Axially Swept Light Sheet Microscopy. Biophys. J. 108, 2807–2815 (2015).

18. Dean, K. M. et al. Isotropic imaging across spatial scales with axially swept light-sheet microscopy. Nat. Protoc. 17, 2025–2053 (2022).

19. Ouyang, Z. et al. Elucidating subcellular architecture and dynamics at isotropic 100-nm resolution with 4Pi-SIM. Nat. Methods 1–13 (2024) doi:10.1038/s41592-024-02515-z.

20. Wang, Q. et al. 4Pi-SIMFLUX: 4Pi single-molecule localization microscopy with structured illumination. Nat. Methods https://doi.org/10.1038/s41592-025-02908-8 (2025) doi:10.1038/s41592-025-02908-8.

21. Aakhte, M. et al. Isotropic, aberration-corrected light sheet microscopy for rapid high-resolution imaging of cleared tissue. Preprint at 10.1101/2025.02.21.639411 (2025).

22. Guo, M. et al. Rapid image deconvolution and multiview fusion for optical microscopy. Nat. Biotechnol. 38, 1337–1346 (2020).

23. Weigert, M. et al. Content-aware image restoration: pushing the limits of fluorescence microscopy. Nat. Methods 15, 1090–1097 (2018).

24. Qu, L. et al. Self-inspired learning for denoising live-cell super-resolution microscopy. Nat. Methods 21, 1895–1908 (2024).

25. Qiao, C. et al. Zero-shot learning enables instant denoising and super-resolution in optical fluorescence microscopy. Nat. Commun. 15, 4180 (2024).

26. Chen, J. et al. Three-dimensional residual channel attention networks denoise and sharpen fluorescence microscopy image volumes. Nat. Methods 18, 678–687 (2021).

27. Li, Y. et al. Incorporating the image formation process into deep learning improves network performance. Nat. Methods 19, 1427–1437 (2022).

28. Qiao, C. et al. Evaluation and development of deep neural networks for image super-resolution in optical microscopy. Nat. Methods 18, 194–202 (2021).

29. Qiao, C. et al. Rationalized deep learning super-resolution microscopy for sustained live imaging of rapid subcellular processes. Nat. Biotechnol. 41, 367–377 (2023).

30. Liu, J. et al. Bio-friendly and high-precision super-resolution imaging through self-supervised reconstruction structured illumination microscopy. Nat. Methods https://doi.org/10.1038/s41592-025-02966-y (2025) doi:10.1038/s41592-025-02966-y.

31. Guo, M. et al. Deep learning-based aberration compensation improves contrast and resolution in fluorescence microscopy. Nat. Commun. 16, 313 (2025).

32. Park, H. et al. Deep learning enables reference-free isotropic super-resolution for volumetric fluorescence microscopy. Nat. Commun. 13, 3297 (2022).

33. Ning, K. et al. Deep self-learning enables fast, high-fidelity isotropic resolution restoration for volumetric fluorescence microscopy. Light Sci. Appl. 12, 204 (2023).

34. Han, J. et al. System- and sample-agnostic isotropic three-dimensional microscopy by weakly physics-informed, domain-shift-resistant axial deblurring. Nat. Commun. 16, 745 (2025).

35. Shocher, A., Cohen, N. & Irani, M. Zero-Shot Super-Resolution Using Deep Internal Learning. in 2018 IEEE/CVF Conference on Computer Vision and Pattern Recognition 3118–3126 (IEEE, Salt Lake City, UT, 2018). doi:10.1109/CVPR.2018.00329.

36. Cheng, Z., Xiong, Z., Chen, C., Liu, D. & Zha, Z.-J. Light Field Super-Resolution with Zero-Shot Learning. in 2021 IEEE/CVF Conference on Computer Vision and Pattern Recognition (CVPR) 10005–10014 (IEEE, Nashville, TN, USA, 2021). doi:10.1109/CVPR46437.2021.00988.

37. Soh, J. W., Cho, S. & Cho, N. I. Meta-Transfer Learning for Zero-Shot Super-Resolution. Preprint at https://doi.org/10.48550/arXiv.2002.12213 (2020).

38. Chen, H. et al. Low-Res Leads the Way: Improving Generalization for Super-Resolution by Self-Supervised Learning. in 2024 IEEE/CVF Conference on Computer Vision and Pattern Recognition (CVPR) 25857–25867 (IEEE, Seattle, WA, USA, 2024). doi:10.1109/CVPR52733.2024.02443.

39. Gustafsson, M. G. L. et al. Three-Dimensional Resolution Doubling in Wide-Field Fluorescence Microscopy by Structured Illumination. Biophys. J. 94, 4957–4970 (2008).

40. Ma, C., Tan, W., He, R. & Yan, B. Pretraining a foundation model for generalizable fluorescence microscopy-based image restoration. Nat. Methods 21, 1558–1567 (2024).

41. Ma, Y. et al. MAGNET: an all-in-one foundation model for cross-modal and cross-dimensional microscopic image restoration. https://doi.org/10.64898/2025.12.23.696141 doi:10.64898/2025.12.23.696141.

42. Qiao, C. et al. Fast-adaptive super-resolution lattice light-sheet microscopy for rapid, long-term, near-isotropic subcellular imaging. Nat. Methods 22, 1059–1069 (2025).

